# Blue light-induced gene expression alterations in cultured neurons are the result of phototoxic interactions with neuronal culture media

**DOI:** 10.1101/783084

**Authors:** Corey G. Duke, Katherine E. Savell, Robert A. Phillips, Jeremy J. Day

**Author notes:** Corresponding author *Correspondence to Jeremy Day* ( | day-lab.org | @DayLabUAB).

## Abstract

Blue waveform light is used as an optical actuator in numerous optogenetic technologies employed in neuronal systems. However, the potential side effects of blue waveform light in neurons has not been thoroughly explored, and recent reports suggest that neuronal exposure to blue light can induce transcriptional alterations *in vitro* and *in vivo*. Here, we examined the effects of blue waveform light in cultured primary rat cortical neurons. Exposure to blue light (470nm) resulted in upregulation of several immediate early genes (IEGs) traditionally used as markers of neuronal activity, including *Fos* and *Fosb*, but did not alter the expression of circadian clock genes *Bmal1*, *Cry1*, *Cry2*, *Clock*, or *Per2*. IEG expression was increased following 4 hours of 5% duty cycle light exposure, and IEG induction was not dependent on light pulse width. Elevated levels of blue light exposure induced a loss of cell viability *in vitro*, suggestive of overt phototoxicity. Changes in gene expression induced by blue waveform light were prevented when neurons were cultured in a photoinert media supplemented with a photostable neuronal supplement instead of commonly utilized neuronal culture media and supplements. Together, these findings suggest that light-induced gene expression alterations observed *in vitro* stem from a phototoxic interaction between commonly used media and neurons, and offer a solution to prevent this toxicity when using photoactivatable technology *in vitro*.

---

OPTICALLY-DRIVEN TECHNOLOGY has been widely adopted in neuroscientific investigation over the past 15 years^1,2^, opening new avenues into experimental design by allowing unprecedented spatial and temporal control over neuronal firing, protein signaling, and gene regulation. Blue waveform light (~470nm) is most often used as the actuator of these technologies. For instance, channelrhodopsin^1^ is a light-gated ion channel that responds to blue light to allow for experimental control over neuronal firing. Similarly, cryptochrome 2 (Cry2)^3–5^ and light-oxygen sensitive protein (LOV) based systems^6–8^ utilize blue light to regulate protein binding and gene expression. Additionally, genetically-encoded calcium sensor technologies to visualize neuronal activity states are becoming more widely utilized both *in vivo* and *in vitro*, and these sensors often rely on prolonged or repeated blue light exposure^9–11^. Together, these optically-driven technologies provide robust experimental control and have enabled new insights into neuronal functioning in healthy and diseased states. However, increased use of these technologies in neuroscience also warrants a more complete understanding of potential off-target effects of prolonged exposure to blue light.

While the phototoxic effects of both ambient and targeted light on cell viability *in vitro* has been noted for decades^12–14^, recent reports documenting blue light-induced gene expression alterations both *in vitro* and *in vivo* have emphasized deleterious effects of blue light on cellular function^15,16^. Multiple reports have documented robust effects of blue light exposure *in vitro*, including upregulation of genes such as *Fos* (aka *cFos*) that are often used as markers of neuronal activity but which can also be induced in response to cellular stress^15–17^. Others have noted that cellular phototoxicity is often the result of reactive oxygen species (ROS) generated in culture media during photostimulation, which can be prevented by utilizing a non-light-reactive media instead of the typical media utilized in neuronal cultures^18^. To our knowledge, it has not yet been determined if the blue light-induced expression alterations of activity-dependent genes observed *in vitro* are the result of a stress response stemming from the culture conditions.

In the present work, we characterized the effects of blue light on gene expression and cell viability *in vitro* using a rat primary neuronal culture model. As recent reports indicate that ROS are generated when culture media is exposed to blue waveform light^14,15^, we hypothesized that light-induced alterations in gene expression would be dependent on the neuronal cell culture media utilized in these experiments. We replicated and extended previous literature by demonstrating that blue light exposure induces multiple IEGs in neuronal cultures, and characterized the duration, frequency, and temporal properties of this effect. Notably, we found that replacing cell culture media with a photostable media supplemented with antioxidants prevented blue light-induced gene expression alterations. Together, these experiments provide insight into the mechanism underlying the unwanted “off-target” effects observed when using optically-driven technology, and offer a path forward to achieving a more precise level of experimental control *in vitro*.

## Results

### Blue light induces immediate early gene expression in cultured primary neurons

To investigate the effects of blue light exposure on gene expression in cultured neurons, we exposed 11 days *in vitro* (DIV) primary cortical cultures to 470nm light and monitored gene expression with reverse transcription quantitative PCR (RT-qPCR; Fig. 1). Cortical neurons cultured in standard media conditions (complete Neurobasal supplemented with B27) were placed on top of a blue LED array light box^22^ inside of a standard cell culture incubator. Pulsed 470nm light was delivered across 7 duty cycle conditions for 0.5 to 8 hrs, followed by RT-qPCR to compare gene expression of light-exposed plates to control plates that were not exposed to light (Fig. 1a). First, neuronal cultures were exposed to 5% duty cycle light for 8 hr, and RNA was extracted to examine the effects of blue light exposure on immediate early gene (IEG) expression. RT-qPCR revealed significant induction of *Fos*, *Fosb*, *Egr1*, and *Arc* mRNA, but not mRNA arising from *Bdnf-IV* (Fig. 1b). To determine if blue light exposure had an effect on the circadian clock, expression of the circadian rhythm genes *Bmal1*, *Clock*, *Per2*, *Cry2*, and *Cry1* was measured under same light exposure conditions. In contrast to robust changes in IEGs, no significant light-induced changes were documented at these key circadian rhythm genes (Fig. 1c).

**Figure 1.**
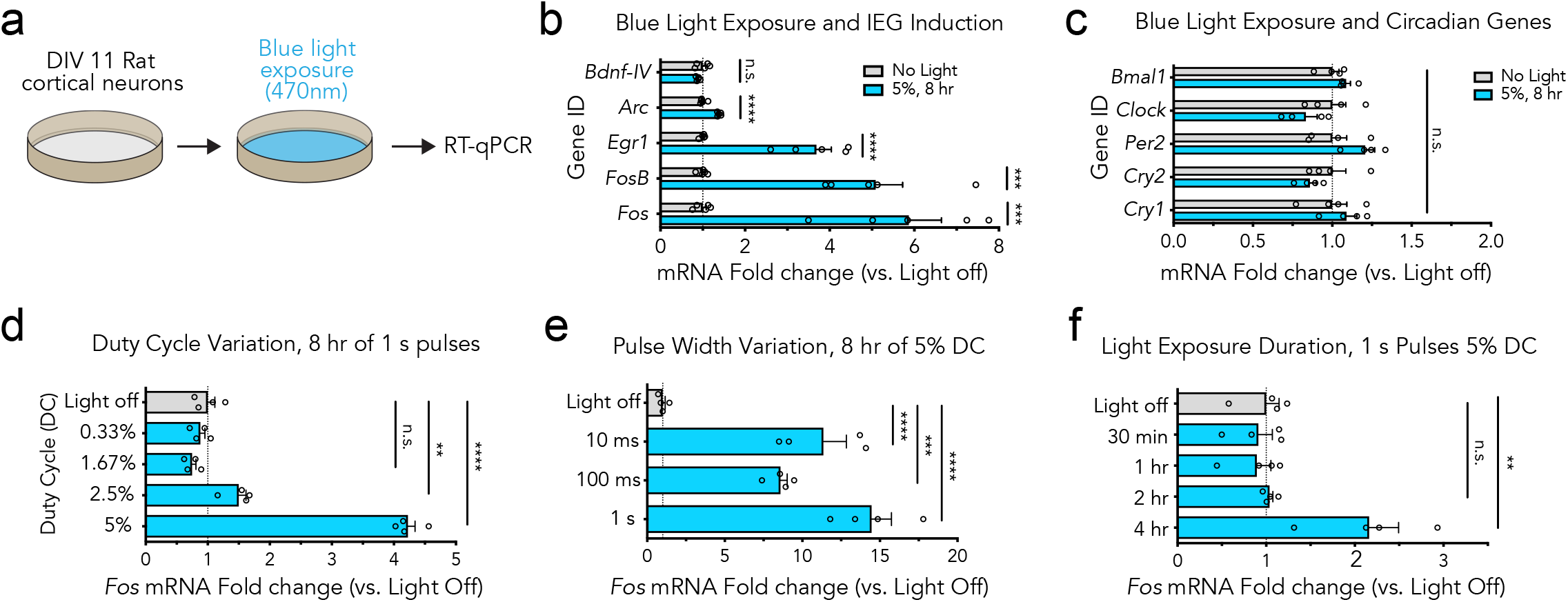
Blue light induces immediate early gene expression in cultured primary neurons. **(a)** Illustration of the experimental design. Primary rat cortical cultures were placed on top of a light box and exposed to blue (470nm) light prior to measurement of gene expression with RT-qPCR. **(b)** Blue light induces gene expression alterations at multiple immediate early genes (n=5, unpaired *t*-test; *Fos t*_(8)_ = 6.301, *P* = 0.0002; *Fosb t*_(8)_ = 6.384, *P* = 0.0002; *Egrl t*_(8)_ = 7.613, *P* < 0.0001; *Arc t*_(8)_ = 10.54, *P* < 0.0001; *Bdnf-IV t*_(8)_ = 1.563, *P* = 0.1566). **(c)**. Circadian rhythm machinery genes were not altered by this blue light exposure (n=4, unpaired *t*-test; *Bmail t*_(6)_ = 1.772, *P* = 0.1268; *Ciock t*_(6)_ = 1.499, *P* = 0.1845 *Per2 t*_(6)_ = 1.910, *P* = 0.1048; *Cry2 t*_(6)_ = 1.491, *P* = 0.1865; *Cryl t*_(6)_ = 0.7978, *P* = .4554). **(d)** *Fos* gene expression alterations are dependent on the amount of light exposure received (n = 4, One-Way ANOVA; F_(4, 15)_ = 215.1, *P* < 0.0001). **(e)** Gene induction is not dependent on pulse width when duty cycle is held constant (n = 4, One-Way ANOVA; F_(3, 12)_ = 32.96, *P* < 0.0001). **(f)** Gene expression is altered as early as 4 hr after light exposure (n = 4, One-Way ANOVA; F_(4, 15)_ = 9.075, *P* = 0.0006). All data are expressed as mean ± s.e.m. Individual comparisons, \*\**P* < 0.01, \*\*\**P* < 0.001, \*\*\*\**P* < 0.0001.

### Blue light is phototoxic to cultured primary neurons

To understand if light-induced gene expression alterations corresponded with changes in cell health, we next examined the effects of blue light exposure on cell viability (Fig. 2). Primary cortical cultures were exposed to blue light (470nm) for 8 hr (at 1.67%, 3.33%, and 6.67% duty cycles) before assessing cell health using fluorescence measurements in a Calcein AMviabilityassay(Fig. 2a-b). We observeddecreased fluorescence intensity at both 3.33% and 6.67% light exposure as compared to a no-light control, indicative of cell death at these duty cycles (Fig. 2c). These findings suggest that cellular health is significantly impacted during sustained light exposure, correlating IEG induction with a loss in cellular viability.

**Figure 2.**
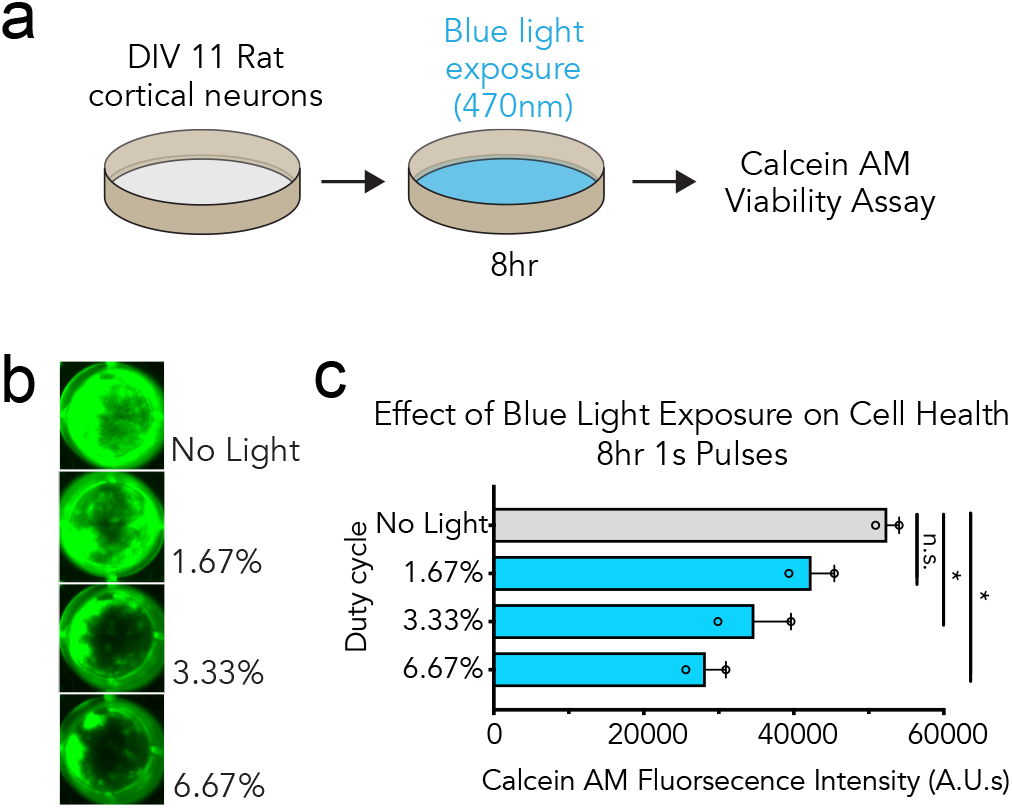
Blue light is phototoxic to cultured primary neurons. **(a)** Illustration of the experimental design. Primary rat cortical cultures were exposed to blue waveform light before cell viability was assessed with a Calcein AM assay. **(b)**. Blue light causes a loss in cell viability with increased exposure. **(c)** Quantified effects of blue light exposure on cell viability at different duty cycles (n = 2, One-Way ANOVA; F_(3, 4)_ = 10.20, *P* = 0.0241). All data are expressed as mean ± s.e.m. Individual comparisons, \**P* < 0.05.

### Photoinert media protects cultured neurons from blue light-induced gene expression alterations

Recent reports suggest light-induced cell viability losses can be overcome with photoinert media^18^, but it remains unclear if light-induced gene expression effects are also dependent on the culture media utilized in these experiments. To examine the contributions of culture media to light-induced gene expression changes, we explored the effects of light exposure in neurons cultured in photoinert media (Fig. 3). Culture media was replaced 12 hr before light exposure with a full or half media change to either Neumo + SOS or Neurobasal + B27 prior to blue light exposure (8 hr at 5% duty cycle) (Fig. 3a). Interestingly, both a full and a half media change to photoinert media completely blocked light-induced *Fos* mRNA increases observed when using standard neuronal culture media (Fig. 3b). To confirm that neurons cultured in photoinert media remained physiologically capable of *Fos* gene induction, we depolarized neurons for 1 hr with potassium chloride (KCl, 25mM) stimulation in this media and observed significant upregulation of *Fos* mRNA (Fig. 3c). Taken together, these results suggest that light-induced upregulation of IEGs in cultured neuron experiments are the result of an interaction with light and culture media, not the result of a direct cellular response to light.

**Figure 3.**
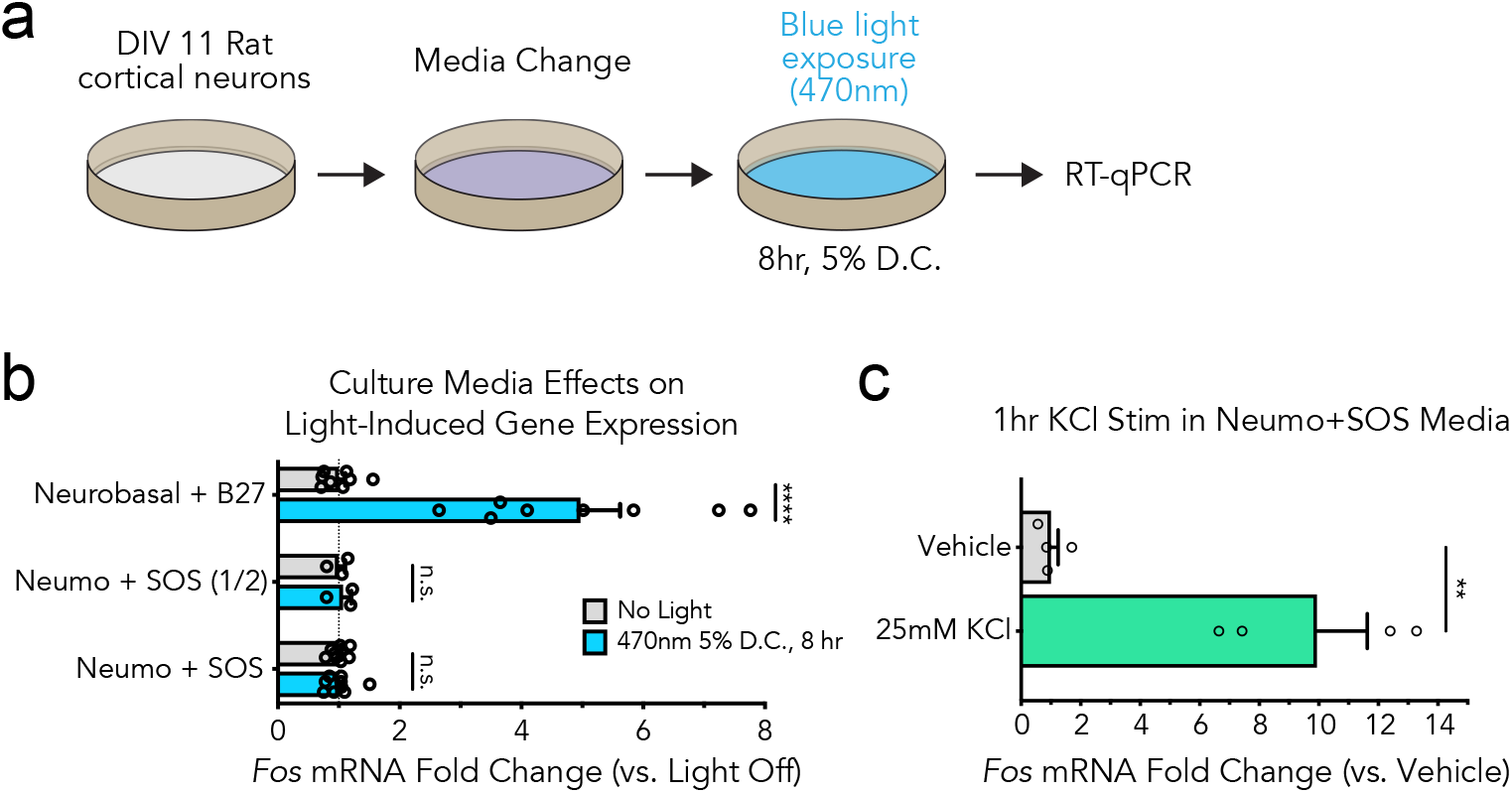
Photoinert media protects cultured neurons from blue light-induced gene expression alterations. **(a)** Illustration of the experimental design. Primary rat cortical cultures were exposed to blue waveform light 12 hr following a media change and then gene expression was assessed by RT-qPCR. **(b)**. Blue light exposure does not induce *Fas* mRNA changes in photoprotective culture media, even if only a half media change is performed (n = 3-9, unpaired *t*-test; Neurobasal *t*_(14)_ = 6.012, *P* = 0.000032; Neumo (1/2) *t*_(4)_ = 0.4099, *P* = 0.708249; Neumo (Full) *t*_(16)_ = 0.02414, *P* = 0.981036). **(c)** *Fas* mRNA can be induced by a 1 hr 25mM KCl stimulation in photoprotective media indicating that the cultures are still capable of induced gene expression alterations (n = 4, unpaired *t*-test, two-tailed; *t*_(6)_ = 5.221, *P* = .0020). All data are expressed as mean ± s.e.m. Individual comparisons, \*\**P* < 0.01, \*\*\*\**P* < 0.0001.

## Discussion

The increased adoption of optical techniques requiring prolonged light exposure in neuroscience highlights a pressing need to both characterize and overcome any off-target effects due to lightexposure alone. To better understand the effects of blue light exposure in cultured neurons, we exposed primary neuronal cultures to blue waveform light and monitored gene expression alterations and cell viability changes. We observed significant elevation of multiple IEGs in primary neuronal cultures in response to blue waveform light, noting that this induction is dependent on the amount of light delivered, and that alterations occur after 4 hr of photostimulation or more. The IEGs we characterized are downstream of the ERK/MAPK pathways and upregulated in response to robust synaptic activation during long term plasticity induction^24–26^. However, these genes are also triggered in response to cellular stress, including exposure to reactive oxygen species^17,27,28^. In contrast, we observed no alterations in expression of circadian rhythm machinery genes, suggesting that this IEG response was not due to light-induced alterations of the circadian cycle. To determine if this transcriptional response is indicative of cellular stress, we examined cell viability across increasing light exposures, demonstrating a decrease in cell viability with increasing amounts of blue light. These results suggest that the gene expression changes we observed following blue light exposure are associated with a cellular stress response.

Previous reports have found that culture media and its supplements can react with light to generate ROS, and recent efforts to overcome this have resulted in the generation of photostable culture media which prevents a decay in cell health during sustained light exposure^12,14,15,18^. Importantly, we report that blue light-induced alterations in IEGs such as *Fos* are prevented when neuronal culture media is transitioned to photostable solution supplemented with antioxidants before light exposure. While in this photostable media, neurons maintain their ability to elicit IEG induction following strong depolarization, indicating that the light-induced gene response is dependent on culture media and can be readily overcome.

With the rapid and widespread adoption of light-inducible technologies in neurobiology^29^, these results provide a path forward when utilizing these techniques *in vitro*. Recent reports have documented light-induced gene expression alterations of *Fos in vivo*^30^, which may be the result of a similar stress response from poor heat dissipation during extended exposure times *in vivo*^31^. In sum, our study highlights the importance of experimental design when using photoactivatable and imaging technologies. Specifically, these results highlight the necessity of including a light exposure only control group when adapting these promising techniques to particular experimental conditions, and the utilization of photostable culture media wherever possible. Improving experimental precision and accuracy is of high priority given the remarkable experimental control and power these techniques provide. Together, the approach outlined here offers an easily implementable solution for the integration of photoactivatable technologies to neuroscientific inquiry *in vitro* that mitigates experimental confounds due to phototoxicity.

## Methods

### Animals

All experiments were performed in accordance with the University of Alabama at Birmingham Institutional Animal Care and Use Committee. Sprague-Dawley timed pregnant rat dams were purchased from Charles River Laboratories. Dams were individually housed until embryonic day 18 for cell culture harvest in an AAALAC-approved animal care facility on a 12-hour light/dark cycle with *ad libitum* food and water.

### Neuronal Cell Cultures

Primary rat neuronal cultures were generated from embryonic day 18 (E18) rat cortical tissue, as described previously^19–21^. Briefly, cell culture plates (Denville Scientific Inc.) were coated overnight with poly-L-lysine (Sigma-Aldrich; 50 µg/ml) and rinsed with diH_2_O. Dissected cortical tissue was incubated with papain (Worthington LK003178) for 25 min at 37°C. After rinsing in complete Neurobasal media (supplemented with B27 and L-glutamine, Invitrogen), a single cell suspension was prepared by sequential trituration through large to small fire-polished Pasteur pipettes and filtered through a 100 µm cell strainer (Fisher Scientific). Cells were pelleted, re-suspended in fresh media, counted, and seeded to a density of 125,000 cells per well on 24-well culture plates (65,000 cells/cm^2^) or 6-well MEA plates (325,000 cells/cm^2^). Cells were grown in complete Neurobasal media for 11 days *in vitro* (DIV 11) in a humidified CO2 (5%) incubator at 37°C with half media changes at DIV 1 and 5. On DIV 10, cells received either a half or full change to complete Neurobasal media, or complete NEUMO media (Cell Guidance Systems; M07-500) supplemented with SOS (Cell Guidance Systems; M09-50) and Glutamax (Thermo Fisher; 35050061), as indicated above.

### Illumination

A custom built 12 LED array was used to illuminate cells, as previously described^22^. Three series of four blue LEDs (Luxeon Rebel Blue (470nm) LEDs; SP-05-B4) regulated by a 700mA BuckPuck (Luxeon STAR) were mounted and soldered onto a rectangular grid circuit board (Radioshack) and positioned inside a plastic enclosure (Radioshack) beneath transparent plexiglass (2cm thick). Cultured neurons were placed atop this enclosure and illuminated from below. An Arduino Uno was used to control LED arrays, delivering light in 1 second pulses at the frequencies required to achieve specifc duty cycles. In all experiments, duty cycle percentage was defined as light on time/total time*100. Aluminum foil was placed on top of the culture dish and enclosure during light delivery. No-light control culture plates were placed atop an identical LED enclosure and wrapped in foil. All handling of culture plates was performed under red light conditions after DIV 5.

### RNA Extraction and RT-qPCR

Total RNA was extracted (RNAeasy kit, Qiagen) and reverse-transcribed (iScript cDNA Synthesis Kit, Bio-Rad). cDNA was subject to RT-qPCR for genes of interest, as described previously^19,23^. A list of PCR primer sequences is provided in **Table 1**.

### Calcein AM Viability Assay

Cell viability was assessed using a Calcein AM Cell Viability Assay Kit (Trevigen; 4892-010-K) according to manufacturer’s instructions for adherent cells. Briefly, cell culture media was removed followed by a wash with 400µl of Calcein AM DW Buffer. 200ul of Calcein AM DW Buffer and 200ul of Calcein AM Working Solution were then added to the culture well and allowed to incubate at 37°C in a humidified CO2 (5%) incubator for 30 min. Culture well florescence was then assessed under 470nm excitation in a standard plate imager (Azure Biosystems c600), and quantified in ImageJ by taking the background subtracted mean pixel value of identical regions of interest areas encompassing individual culture wells.

### Statistical Analysis

Transcriptional differences from RT-qPCR experiments were compared with either an unpaired *t*-test or one-way ANOVA with Dunnett’s or Tukey’s *post-hoc* tests where appropriate. Statistical significance was designated at α = 0.05 for all analyses. Statistical and graphical analyses were performed with Prism software (GraphPad). Statistical assumptions (e.g., normality and homogeneity for parametric tests) were formally tested and examined via boxplots.

### Data Availability

All relevant data that support the findings of this study are available by request from the corresponding author (J.J.D.).

## Acknowledgements

This work was supported by NIH grants DA039650, DA034681, and MH114990 (J.J.D.), NS061788 (C.G.D.), DA042514 (K.E.S.). Additional assistance to J.J.D. was provided by the UAB Pittman Scholars Program.

## Author contributions

C.G.D., K.E.S., and J.J.D conceived of the experiments. C.G.D. and K.E.S. performed the experiments with assistance from R.A.P. C.G.D., K.E.S., and R.A.P. performed statistical and graphical analysis, and data interpretation. C.G.D. and J.J.D. wrote the manuscript. J.J.D. supervised all work. All authors have approved the final version of the manuscript.

## Competing Interests

The authors declare no competing financial interests.

